# Exploring the miRNA-mediated response to combined stresses in melon plants

**DOI:** 10.1101/2021.07.30.454429

**Authors:** Pascual Villalba-Bermell, Joan Marquez-Molins, María-Carmen Marques, Andrea G. Hernandez-Azurdia, Julia Corell-Sierra, Belén Picó, Antonio J. Monforte, Santiago F. Elena, Gustavo G. Gomez

## Abstract

Climate change has been associated with a higher incidence of combined adverse environmental conditions that can promote a significant decrease in crop productivity. However, knowledge on how a combination of stresses might affect plant development is still scarce. MicroRNAs (miRNAs) have been proposed as potential targets for improving crop-productivity. Here, we have combined deep-sequencing, computational characterization of responsive miRNAs and validation of their regulatory role in a comprehensive analysis of melon’s response to several combinations of four stresses (cold, salinity, short day, and infection with a fungus). Twenty-two miRNA families responding to double and/or triple stresses were identified. The regulatory role of the differentially expressed miRNAs was validated by quantitative measurements of the expression of the corresponding target genes. A high proportion (ca. 60%) of these families (mainly highly conserved miRNAs targeting transcription factors) showed a non-additive response to multiple stresses in comparison with that observed under each one of the stresses individually. Among those miRNAs showing non-additive response to stress-combinations, most interactions were negative suggesting the existence of functional convergence in the miRNA-mediated response to combined stresses. Taken together, our results provide compelling evidences that the response to combined stresses cannot be easily predicted from the study individual stresses.

## 1 INTRODUCTION

During their life cycle, plants are exposed to a wide array of adverse environmental conditions that, in general, limit their normal development and productivity. These complex interactions result in several stress situations that disturb the cell’s homeostasis negatively affecting plant-growth. Consequently, stress-induced damages in productivity are the primary cause of extensive agricultural losses worldwide (Priya et al., 2019). Reduction in crop yield due to environmental variations has increased steadily over the last decades. In addition, several production models project a reduction in the yields of major agricultural crops in the future, mostly due to climatic changes (Rosenzweig et al., 2014).

Climate change, entailing shifts in temperature, precipitation and atmospheric composition, among other factors, represents a moving target for plant developmental adaptation. In parallel, environmental modifications can favor the development of new plant-pest and/or pathogens or increase the incidence levels of already existing ones. As a consequence of this complex environmental scenario, it is expected that combined abiotic and biotic stresses can affect plants at the level of molecular functions, developmental processes, morphological traits, and physiology, resulting in a significant decrease in crop production and quality (Gray & Brady, 2016; Morales-Castilla et al., 2020).

Multiples studies focused on plant responses to individual stresses have been carried out over the last years. However less attention has been paid to the effect that combinations of adverse environmental conditions might exert on plant development. In order to improve crop yield and to meet the growing challenges stemming from rapid population growth, extensive efforts are needed to understand the mechanisms underlying plant responses to simultaneous exposure to multiple stresses (Zhang & Sonnewald, 2017). Previous works have pointed out that studying stress conditions separately would not allow to infer the expected plant response to multiple stresses. Using *Arabidopsis thaliana* as experimental model, it was shown that the response to a combination of drought and heat was unique and could not be directly extrapolated from the plant response to each stress applied individually (Rizhsky et al., 2004; Suzuki et al., 2005; Rossel et al., 2007). Similar findings were also reported for a combination of heat and high light intensity in sunflower (Hewezi, Léger & Gentzbittel, 2008), and heat and salinity in wheat (Keleş & Öncel, 2002). Consequently, plant response to combined adverse environmental conditions should be handled as a new state of stress that requires a novel conceptual viewpoint (Mittler & Blumwald, 2010).

In general, plants respond to stress conditions through a complex reprogramming of their transcriptional activities aiming to reduce the impact of stress on their physiological and cell homeostasis. Environmental variations have selected diverse responses among plant lineages, landraces and wild crops relatives. Studies on natural variations can provide novel insights into evolutionary processes modulating stress response (Meyers et al., 2008; Haak et al., 2017). Elucidation of how endogenous regulators and the environment interact during plant development is a long-standing grand challenge in modern biology as well as in crop breeding (Lovell et al., 2015).

MicroRNAs (miRNAs) play a versatile role as regulators of gene expression. Plant genes encoding miRNAs are transcribed by RNA polymerase II as primary transcripts harboring a fold back structure that is processed by DICER-LIKE 1 (DCL1) in a duplex (21 or 22 nt in length) which once 2’-O-methylated by HEN1 is loaded into an AGO complex (Bartel, 2004; Bologna & Voinnet, 2014; Reis, Eamens & Waterhouse, 2015; Achkar, Cambiagno & Manavella, 2016). miRNAs regulate gene expression by means of sequences complementarity with both RNA and DNA targets (Song, Li, Cao & Qi, 2019). Their functions include modulation of a vast array of plant biological processes related to grown and development (Bologna & Voinnet, 2014), including the recovering of the plant-cell homeostasis during exposure to adverse environmental condition (Song et al., 2019; Xu et al., 2019). In addition, it has been recently described that the biogenesis and turnover of certain miRNAs is also susceptible to be controlled by external stimulus (Bustamante et al., 2018; Manavella, Yang & Palatnik, 2019). Indeed, it has been proposed that miRNAs are ideal targets to be manipulated to improve crop productivity (Tang & Chu, 2017; Xu et al., 2019). However, most of the described stress-responsive miRNAs come from rice and tomato, as very few miRNAs have been investigated in detail in other crops. Henceforth, additional efforts are needed to decipher the role of miRNA-mediated responses to adverse environmental conditions in other economically relevant crops (Tang & Chu, 2017).

Although, increasing evidences support the role of miRNAs as key modulators of plant response to both biotic (Sun, Niu & Fan, 2017; Xie et al., 2017; Brant & Budak, 2018) and abiotic stress conditions (Cervera-Seco et al., 2019; Wang et al., 2020; Cheng et al., 2021; Zhao et al., 2021), research focusing on elucidating the regulatory role of the miRNAs during exposure to combined adverse environmental conditions is still scarce (Xu et al., 2019) and only a few studies considering the effects of an unique combination of stresses have been addressed in soybean (Ning et al., 2019) and *A. thaliana* (Gupta, Patil, Qamar & Senthil-Kumar, 2020).

Melon (*Cucumis melo*) is one of the cucurbit crops with more economic impact. Melon has a high adaptability to warm and dry climates, so it can be a target crop to cope with the climate change threats. Previous genetic studies in cucurbits have been focused mainly in fruit quality and disease resistance (Gonzalo & Monforte 2017). However, the study of the response to combined stress conditions have not been thoroughly addressed in cucurbits. Consequently, there is a lack of consensus protocols, target traits and, therefore, identification of tolerant genotypes to develop efficiently resilient cultivars.

Here, we use deep-sequencing, computational approaches and specific miRNA-targets quantification to present a comprehensive functional analysis of miRNA expression profiles in response to one triple (cold, salinity and short day) and five double (cold and drought, cold and salinity, cold and short day, drought and salinity, and drought and infection with the fungus *Monosporascus cannonballus*) combinations of stress conditions in melon (*Cucumis melo*), a crop extensively cultivated in semi-arid regions worldwide. The analyzed stress conditions were coincident, in part, with those employed recently to infer the miRNA-mediated regulatory network of response to individual stresses in melon (Sanz-Carbonell et al., 2019; Sanz-Carbonell, Marques, Martinez & Gomez, 2020). The parallelism between both experimental approaches made possible to unambiguously analyze the effects that the combined adverse environmental conditions have on the accumulation of the stress-responsive miRNAs.

## 2 METHODS

### 2.1 Plant material, growth conditions, and stress treatments

Melon seeds of cv. Piel de Sapo were germinated in Petri dishes at 37 °C/48 h in darkness followed by 24 h/25 °C (16/8 light/darkness). Melon seedlings were sown in pots and maintained for 10 days under controlled conditions (28 °C/16 h light and 20 °C/8 h darkness). At day 11, plants were exposed to six stress-combined treatments (detailed in Table S1). At eleven days post-treatment, the first leaf under the apical end per plant was collected in liquid nitrogen and maintained at –80 °C until processing. Each analyzed sample corresponds to a pool of three treated plants. Three biological replicates were performed per treatment. Leaves recovered from non-treated plants were considered as controls.

### 2.2 RNA extraction and small RNA (sRNA) purification and sequencing

Total RNA was extracted from leaves (∼0.1 g) recovered from treated and control melon as previously described (Sanz-Carbonell et al., 2019; Sanz-Carbonell, Marques, Martinez & Gomez, 2020). The low-molecular weight RNA (< 200 nt) fraction was enriched from total RNA using TOTAL-miRNA (miRNA isolation Kit, REAL) according to the manufacturer’s instructions. Production and sequencing of the libraries were carried out by Novogene (https://en.novogene.com). Eighteen cDNA libraries were obtained by following Illumina’s recommendations and sequenced in a HiSeq 2000 (Illumina) equipment. Adaptors and low-quality reads were trimmed by using the cutadapt software. For the sake of comparing the results generated in here with those obtained for single stresses, data previously obtained from melon plants exposed to identical single stress conditions for 11 days (Sanz-Carbonell et al., 2019) were also included in the study. Melon miRNA sequences used in this study have been submitted to the genomic repository SRA of the NCBI and are available in the BioProject (PRJNA741881).

### 2.3 RT-qPCR assays

To analyze the expression of target genes, total RNA (1.5 μg) was subjected to DNase treatment (EN0525, Thermo Scientific™) followed by reverse transcription using RevertAid First Strand cDNA Synthesis Kit (Thermo Scientific™) according to the manufactureŕs instructions for use with oligo-dT. cDNAs were amplified by conventional end-point RT-PCR using specific primers to assess for sequence specificity. Then, real-time PCR was performed as described previously (Bustamante et al., 2018). All analyses were done in triplicate on a QuantStudio qPCR instrument (Thermo Scientific™) using a standard protocol. The efficiency of PCR amplification was derived from a standard curve generated by four 10-fold serial dilution points of cDNA obtained from a mix of all the samples. Relative RNA expression was quantified by the comparative ΔΔ*C_T_* method (Livak & Schmittgen, 2001) and normalized to the geometric mean of Profilin (NM_001297545.1) expression. The statistical significance of the observed differences was evaluated by the paired *t*-test. All primers used were described previously (Sanz-Carbonell et al., 2019).

### 2.4 Bioinformatic analysis of miRNA sequences

To study the correlation exhibited by the miRNA expression profiles among the different stresses and their biological replicates, principal component analysis (PCA) was used. PCA was performed using the prcomp function with scaling in the stats R package v. 4.0.4 (R Core Team 2013). Mann-Whitney-Wilcoxon tests were performed to assess for significant differences in the data clusters for Euclidean distances calculated between groups and among groups with the wilcox.test function in the stats R package.

Differential expression of melon sRNAs was estimated using three R packages NOISeq (Tarazona et al., 2015), DESeq2 (Love, Huber & Anders, 2014) and edgeR (Robinson & Oshlack, 2010) for pairwise differential expression analysis of expression data. Differentially expressed sRNAs were filtered out using three criteria: (i) log_2_-fold change |log_2_*FC*| ≥ 1.25, (ii) adjusted *p* ≤ 0.05 (DESeq2 and edgeR) and probability ≥ 0.95 (NOISeq), and (iii) RPMs ≥ 5 for at least three libraries in control samples or at least two libraries in any stress. sRNAs identified as responding to stress by the three methods were aligned against miRNA sequences in miRBase (release 22) (Kozomara, Birgaoanu & Griffiths-Jones, 2019). Only fully homologous miRNAs to previously described mature melon miRNAs and known *Viridiplantae* miRNAs were kept. Afterwards, these sequences were re-annotated by aligning them against miRNA precursors of melon deposited in miRBase and were considered as known stress-responsive miRNAs. Unaligned sequences were realigned allowing for one mismatch against the melon genome to identify potential precursors. These sequences were also identified as known stress-responsive miRNAs; the rest were discarded. The entire pipeline is shown in Figure S2.

To determine the general sense of the expression for each miRNA family we employed the median value of expression estimated by box-plot analysis of all family-related sequences under each stress condition considering the log_2_*FC* values obtained by edgeR. The most frequent sequence in each miRNA family and stress were used to generate heatmaps with an R interface to morpheus.js heatmap widget (https://github.com/cmap/morpheus.R).

### 2.5 Analysis of the stress combination effect

The expression of reactive miRNAs in response to combined stress conditions can be enfolded in at least one of the three following categories: (i) additive if the observed response to combined stresses is just the sum of the magnitude responses observed for each individual stress, i.e., this represents the null hypothesis of independent actions, (ii) negative if the observed response is smaller than the expected additive response and (iii) positive if the observed value is greater than the expected additive response. In this framework, if a given miRNA shows an additive response upon exposure to two stresses, it can be assumed that both stresses trigger independent miRNA-mediated responses. In contrast, a miRNA showing a significantly negative or positive deviation from the null hypothesis, shall be taken as indicative of a specific response to the combined stresses beyond the simple additive case. To quantitatively test the null hypothesis of additive effects on miRNA-mediated response to stress combinations, we define an *stress combination effect* (*SCE*) index that refers to the miRNA response value to combined stresses in comparison to what should be expected from individual stress conditions as *SCE* = (*C* + *S_ab_*) − (*S_a_* + *S_b_*), where *C* refers to the means of the normalized reads recovered in control, *S_ab_* to the reads observed in plants exposed to combined stresses *a* and *b* and *S_a_* and *S_b_* to the reads arising from each individual stress (Table S6A and Table S6B). For the triple stress condition (*S_abc_*) and additional value (*S_c_*) -referred to the means of normalized reads in the additional stress condition *c*-should be added to the second terms of the equation. Only *SCE* values with a significant false discovery rate (FDR)-adjusted *p* value were considered as reliable indicators of effects of stress-combinations onto miRNA accumulation.

Reads exhibiting zero means values in any of the analyzed combinations were filtered out. The data associated to the miRNA expression under single stress conditions were extracted from a previous work analyzing the differential expression of melon miRNAs in response to seven biotic and abiotic single stress conditions (Sanz-Carbonell *et al*. 2019). The statistical significance of these effects was calculated on the basis of a standard Normal distribution. Then, the 22 stress-responsive miRNA-families were organized in a binary table of presence and absence (Table S7), in which the values one and zero represent, respectively, whether or not a miRNA family has at least a member exhibiting a significant non-additive (positive or negative) effect in response to a combined stress condition. The hclust function in stats R package (v. 4.0.4) was used to compute a hierarchical clustering (HC) specifying Ward linkage (ward.D) as an agglomeration method and using the simple matching coefficient metric to calculate the distance matrix. The statistical significance of the HC was estimated with a Mann-Whitney-Wilcoxon test.

## 3 RESULTS

### 3.1 Stress combinations and sRNAs dataset

High-throughput sequencing of sRNAs was performed starting from 22 (three replicates for each stress condition plus four non-treated controls) sRNA libraries constructed with RNA extracted from leaves of melon plants 11 days after exposure to six (five double and one triple) combined stress conditions: (i) cold and drought (C-D), (ii) cold and salinity (C-Sal), (iii) cold and short day (C-SD), (iv) drought and salinity (D-Sal), (v) drought and *M. cannonballus* infection (D-Mon), and (vi) cold, salinity and short day (C-Sal-SD) (Table S1). Regarding the stress conditions analyzed, we selected abiotic conditions well established as crucial for melon plant development (cold, drought, salinity, and short day) and infection with *M. cannonballus*, a soil-borne fungal pathogen causing root rot and wilting in melon (Pollack & Uecker, 1974). Only sequences with size ranging between 20 - 25 nt in length and nonmatching to rRNA, tRNA, snoRNA, and snRNA sequences deposited in the Rfam data base (http://rfam.xfam.org) were further included in this study. A total of 80,620,994 reads (representing 36,836,230 unique sequences) were recovered. The distribution of reads by stress condition is detailed in Table S2.

Associations between sRNA expression profiles (considering the different treatments and their biological replicates) were evaluated using PCA. The percentages of variance explained by the first three PCs were 20.4%, 17.1% and 13.8%, respectively (adding up to 51.3% of the total observed variance). The PCA plot in Figure 1A shows that biological replicates clustered together (attesting for the reproducibility of our assays) and treatments clearly separated in the PC space with high significance (*p* = 5.886×10^−15^). The sRNAs exhibited a distribution of read lengths strongly enriched for 24 nt long (45.7%), followed by similar accumulations of 21 (13.5%), 22 (12.6%) and 23 (13.5%) nt long molecules. As expected, reads of 20 and 25 nt represented the less abundant categories (5.9% and 8.5%, respectively) (Figure 1B). These differences in accumulation of different sRNA lengths was statistically significant (2-ways non-parametric ANOVA, Table S3:a *p* < 10^−5^). The effect was entirely due to the large enrichment in 24 nt long sRNAs (Dunn’s post hoc pairwise tests, Table S3b: *p* ≤ 0.0134 in all pairwise comparisons) and consistent with what has been previously described in melon (Sattar et al., 2012; Herranz, Navarro, Sommen & Pallas 2015; Sanz-Carbonell et al., 2019; Sanz-Carbonell, Marques, Martinez & Gomez, 2020) and other members of the *Cucurbitaceae* family (Jagadeeswaran et al., 2012). Non-significant differences were found between stress conditions regarding the observed distribution of sRNAs sizes (Table S3a: *p* = 0.857), nor the interaction between both factors (Table S3a: *p* = 0.750).

**Figure 1.**
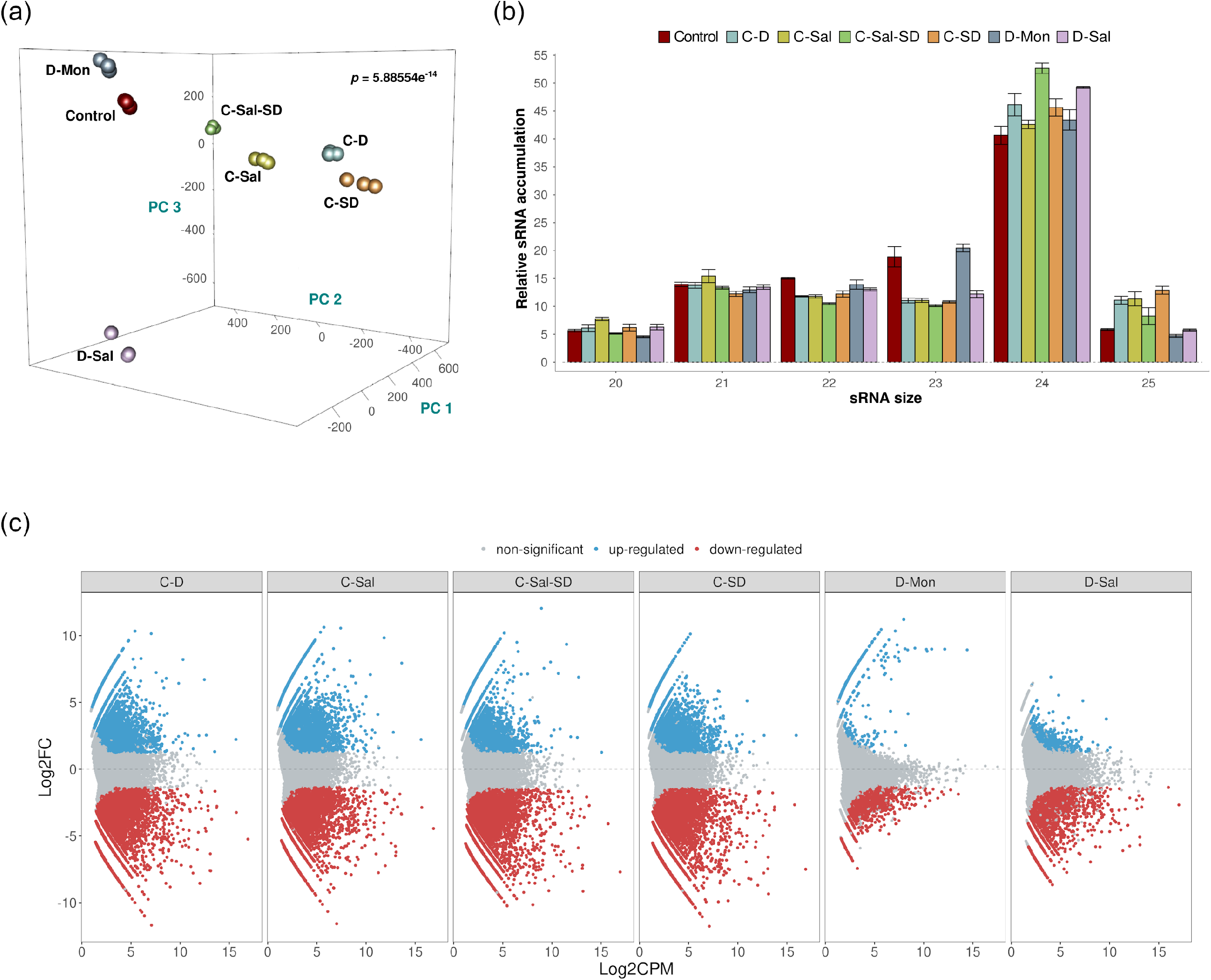
Analysis of the sRNA populations. (a) PCA based on sRNAs accumulation in three biological replicates of melon plants exposed to the six stress combined treatments and controls. The statistical significance (*p* = 5.886×10^−14^) was estimated by Mann-Whitney-Wilcoxon test, considering the inter- and intra-group Euclidean distances. (b) Diagram showing the relative accumulation (and distribution of the total clean reads of melon sRNAs ranging between 20 - 25 nt obtained from the analyzed sequenced libraries. The control and the different analyzed treatments are represented with colors. The shown values represent the sum of all repetitions. Bars indicate the standard error. (c) Graphic representation of the expression values (estimated by edgeR) of sRNA sequences recovered from melon exposed to different stress conditions. The dots indicate the expression value of each sRNA. Red and blue dots indicate significant values for differential expression with |log_2_*FC*| ≥ 1.25, respectively. Grey dots indicate sRNAs with non-significant differential expression.

The effect of the stress conditions onto sRNAs accumulation was evaluated by pairwise comparisons between control and treated samples. As described in section 2.4 above, only sequences that match the conditions |log_2_*FC*| ≥ 1.25 and *p* < 0.05, were considered as significantly differentially expressed and retained for subsequent analysis (Figure S1). A total of 35,906 unique reads fulfilled these conditions. The combinations that included cold as one of the stressors showed the most drastic alteration in sRNAs accumulation (21,592 reactive sRNAs in C-D, 20,760 in C-Sal, 23,506 in C-SD and 21,263 in C-Sal-SD). In contrast, only 1595 and 3988 differentially expressed sRNAs were identified in plants treated with the combination D-Mon and D-S, respectively (Figure S2B). These results support the notion that exposition to low temperature (in any combination) is the most stressful environmental condition, resulting in the strongest alteration of the sRNA metabolism in melon (Figure 1C).

### 3.2 Combined stresses induce a general decrease of miRNA expression

To identify melon miRNAs reactive to combined stress conditions, differentially expressed sRNAs were aligned against miRNA sequences (both mature and precursors) recovered from miRBase (http://www.mirbase.org/). Only sRNAs ranging 20 - 22 nt and fully homologous to database sequences, were considered. Two sequences homologous to mature miR6478 but lacking a known transcript in melon with a canonical hairpin were excluded for subsequent analysis (Figure S1). After filtering, 100 unique sequences belonging to 22 known miRNA families were identified as responsive to the combined stress conditions studied (Table S3). In general, all family-related sequences showed a comparable trend of accumulation in response to the stress conditions analyzed (Figure 2A). A sequence-variant of miR398b (down-regulated in C-D treatment, but showing a minority accumulation rate respect to predominant family-related sequences) and the non-canonical miRNAs derived of the alternative processing of miR319 (miR319nc) (Bustamante et al., 2018) and miR159 (miR159nc) (Bologna, Mateos, Bresso & Palatnik, 2009) precursors (up-regulated in cold-containing combinations and without regulatory activity described yet) showed a discordant response with the family-wise trend. In these two circumstances, the response trend of the more representative family members was considered for ulterior analysis.

**Figure 2.**
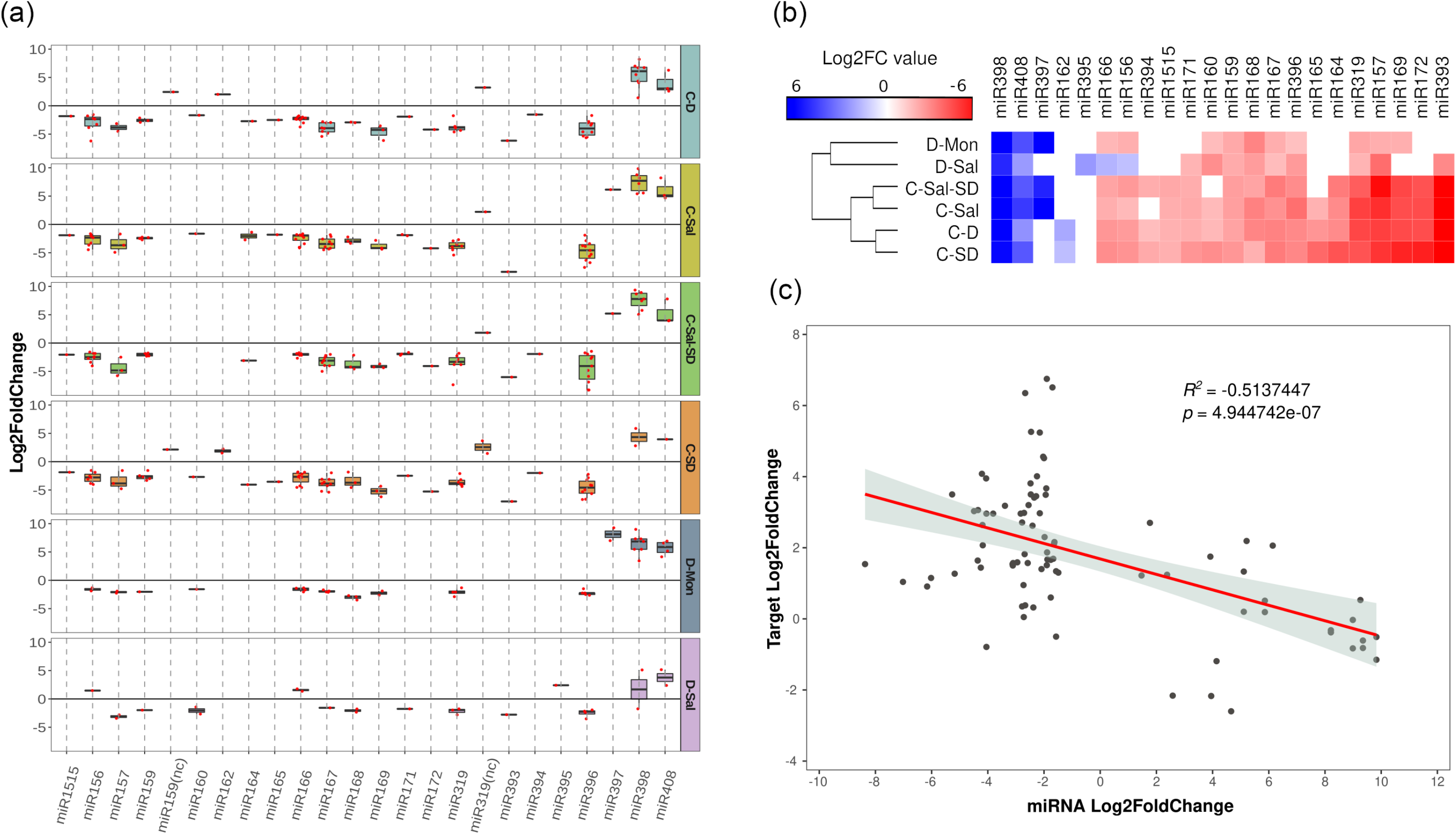
General description of stress-responsive miRNA families. (a) Boxplot analysis showing the general expression value observed for each miRNA-family member. To determine the general sense of the expression for each miRNA family we employed the median value of expression (represented by internal box-line) estimated by boxplot analysis of all family-related sequences. The differential expression values represented in the figure correspond to the log_2_*FC* obtained using edgeR. (b) Heatmap of 22 miRNAs differentially expressed in melon in response to combined stress. The differential expression values represented correspond to the median of the log_2_*FC* values obtained using edgeR for each miRNA family. (c) Scatter plot showing the significant negative correlation (estimated by Pearson correlation coefficient) between the expression levels of 16 selected stress-responsive miRNAs with differential accumulation determined by sequencing and the accumulation of their targets in the corresponding stress conditions, estimated by RT-qPCR.

The general response to stress conditions was the down-regulation of miRNAs (Figure 2B). Sequences included in miRNA families miR157, miR159, miR167, miR168, miR319, and miR396 showed significantly decreased accumulation in all the stress conditions analyzed. Diminished accumulation in response to stress was also observed for miR156, miR160 (except under C-Sal-SD), miR164, miR166, miR169 (except for D-Sal), miR171, miR172 (except for D-Sal and D-Mon), miR393 (except for D-Mon), miR394, and miR1515. Finally, miR165 was down-regulated in three stress conditions involving cold (C-SD, C-D and C-Sal). Regarding miRNAs up-regulated in response to stress, the miR398 and miR408 family-related members (except for the reads related to miR398b described above) showed increased accumulation in all stress conditions, whereas miR159 was significantly overexpressed in response to C-SD and C-D and miR397 family was so in plants exposed to C-Sal, C-Sal-SD and D-Mon. Sequences related to miR156, miR166 and miR395 were specifically up-regulated under D-Sal stress.

The analysis of the miRNA expression focused on each particular stress combination evidenced that cold was the most adverse environmental condition with major impact on miRNA expression in melon. A total of 20 miRNA families were reactive to C-SD and C-D and 19 to C-Sal (Figure 2B and Table S4). While 18 miRNAs families showed differential expression under the combination of three stresses. A weaker response was associated to treatments with D-Sal (14 reactive miRNA families) and D-Mon (13 miRNAs with altered expression). Considering both stress condition and miRNA expression-trend, except miR156 and miR166 (up-regulated in D-Sal and down regulated in the other stress conditions), all miRNAs exhibit a homogenous response to the six combinations of adverse environmental conditions analyzed.

It has been recently proposed that certain melon miRNAs are predominantly reactive to diverse biotic and abiotic stress conditions, while other specifically respond to certain stressor and/or expositions time (Sanz-Carbonell, Marques, Martinez & Gomez, 2020). Based on this particular behavior miRNAs belongings to both different groups were identified as stress responsive miRNAs with *broad* and *narrow* response range, respectively, while a third group that exhibit a moderated reactivity in response to stress were identified as *intermediates*. According to our data, ten miRNA families showed the higher response rate to combined stress, with significant differential expression (either up or down) in the six analyzed conditions (Table S4). Eight of these miRNA families (miR156, miR157, miR166, miR167, miR319, miR396, miR398, and miR408) were mostly coincident with melon miRNAs families classified in the broad response category (*generalists*), while miR159 and miR168 were previously categorized as intermediates. In contrast, miRNAs with a lower response rate to double and triple stresses (responsive in three o less conditions), pervasively pertained to miRNAs families previously reported as showing *specific* response to stress conditions in melon.

To test the functional role of the miRNAs reactive to combined stresses, we analyzed the correlation between miRNA levels and transcripts accumulation in 16 representative miRNA-target modules (Table S5) previously established and validated to occur in melon plants (Bustamante et al., 2018; Sanz-Carbonell et al., 2019; Sanz-Carbonell, Marques, Martinez & Gomez, 2020). We focused on the miRNAs reactive to at least three different stress conditions (miR156, miR159, miR160, miR164, miR166, miR167, miR169, miR171, miR172, miR319, miR393, miR396, miR397, miR398, and miR408). As expected, a significant negative correlation (*r* = −0.514, 83 df, *p* = 4.945×10^−7^) was obtained when the expression values of stress-responsive miRNAs were compared with the accumulation (estimated by RT-qPCR) of their target-transcripts (Figure 2C).

### 3.3 The miRNA-mediated response to stress combinations cannot be predicted from the response to single stresses

To determine the dynamic of the miRNA-mediated response to multiple stress conditions we compare the accumulation levels of stress-responsive miRNAs in plants subjected to the individual stress conditions with those of plants exposed to combined stresses. To do so, we computed *SCE* as defined in section 2.5 above. Except for the combination C-Sal-SD, the additive effect was predominant in number of unique miRNA sequences in the analyzed stress combinations (65.26% of the unique reads) (Figure 3A). However, considering the entire miRNAs population (total reads) a comparable abundance of additive (50.07%) and non-additive (49.93%) instances was observed in response to combined stresses. Interestingly, when evaluating only by miRNA family, 57.58% had at least a member showing a significant (negative or positive) *SCE* value (Figure 3A and Table S7).

**Figure 3.**
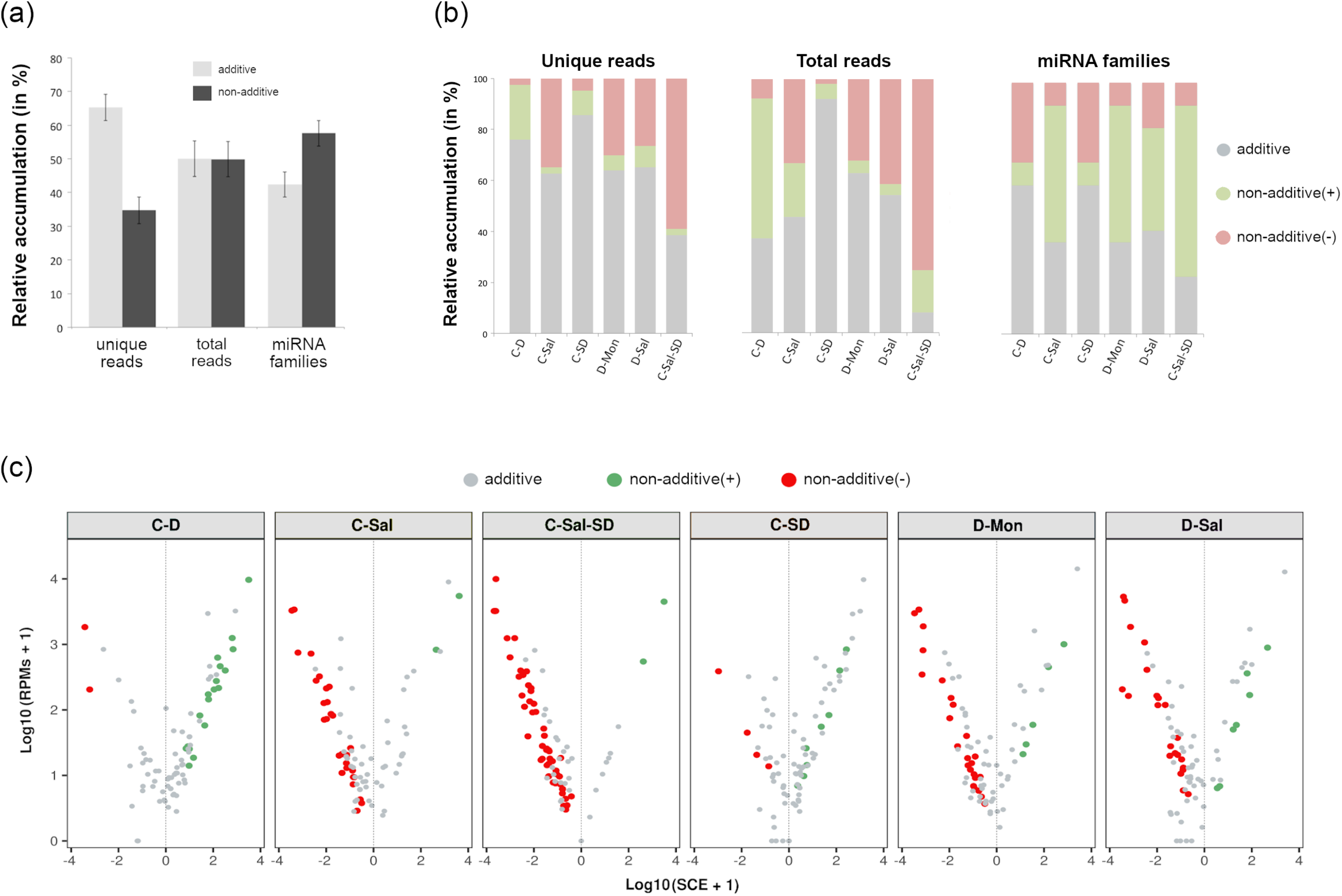
Effects of the stresses combination onto the accumulation rate of stress responsive miRNAs. (a) Graphic representation of the mean percentage for the six analyzed treatments of miRNA related reads that exhibit additive (grey) or non-additive (black) response to combined stress conditions in comparison to single stresses considering unique reads (left columns), total reads (central columns) and miRNA families (right). Bars represent the standard error between means. (b) Detail of the global response rate in each stress condition considering the two (positive or negative) type of possible non-additive response to combined stresses. (c) Volcano plot showing significant positive (green dots) and negative (red dots) *SCE* values obtained for each miRNA-related read, in response to each combined stress condition. miRNAs with non-significant deviations from the additive null model are in grey. More detailed information is provided in the Table S6B.

Regarding significant non-additive interactions, the stress combination predominantly exerted a negative effect in four (C-Sal, D-Sal, D-Mon, and C-Sal-SD) of the six analyzed treatments (Figure 3B). By contrast, in C-D and C-SD, *SCE* > 0 values were the most common. Analyzing each stress combination individually, C-SD was the condition in which miRNAs shown the smallest fraction of specific response to combined stresses (14.46% of unique reads, 7.77% of total reads and 40.91% of the miRNA families). In contrast, a higher differential interaction (76.47% for negative and 2.94% for positive) was observed in response to the triple combinations C-Sal-SD (61.45% of unique reads, 92.05% of total reads and 77.27% of the miRNA families) (Figure 3B). A more general view of the additive and non-additive effects of the combined stresses onto the global population of miRNA-related reads in each analyzed stress condition is showed in the Figure 3C.

Considering the response trend of miRNA family members, we observed that, in general, reads showed a coordinated interaction (*SCE* positive or negative) in response to the combination of stresses (Figure 4A). Consequently, a negative response was also pervasive under a global miRNA-family viewpoint. Exceptions to this rule were observed for the families miR157 in C-SD and miR159 in D-Sal, that contained members showing both positive and negative *SCE* values under the indicated stress combination. However, it is worth nothing that the miRNA sequences with a non-coincident trend are minority relative to the other family members (Table S6A). Therefore, in these two specific cases the response trend of the predominant reads was considered as representative of the family behavior for ulterior analysis (Figure 4B). The highest number (17) of miRNA families showing significant *SCE* values was observed in plants exposed to the triple combinations of stresses, followed by C-Sal and D-Mon (14) and D-Sal (13). In contrast, only nine miRNA-families were identified as significantly interactive in response to C-D and C-SD, respectively.

**Figure 4.**
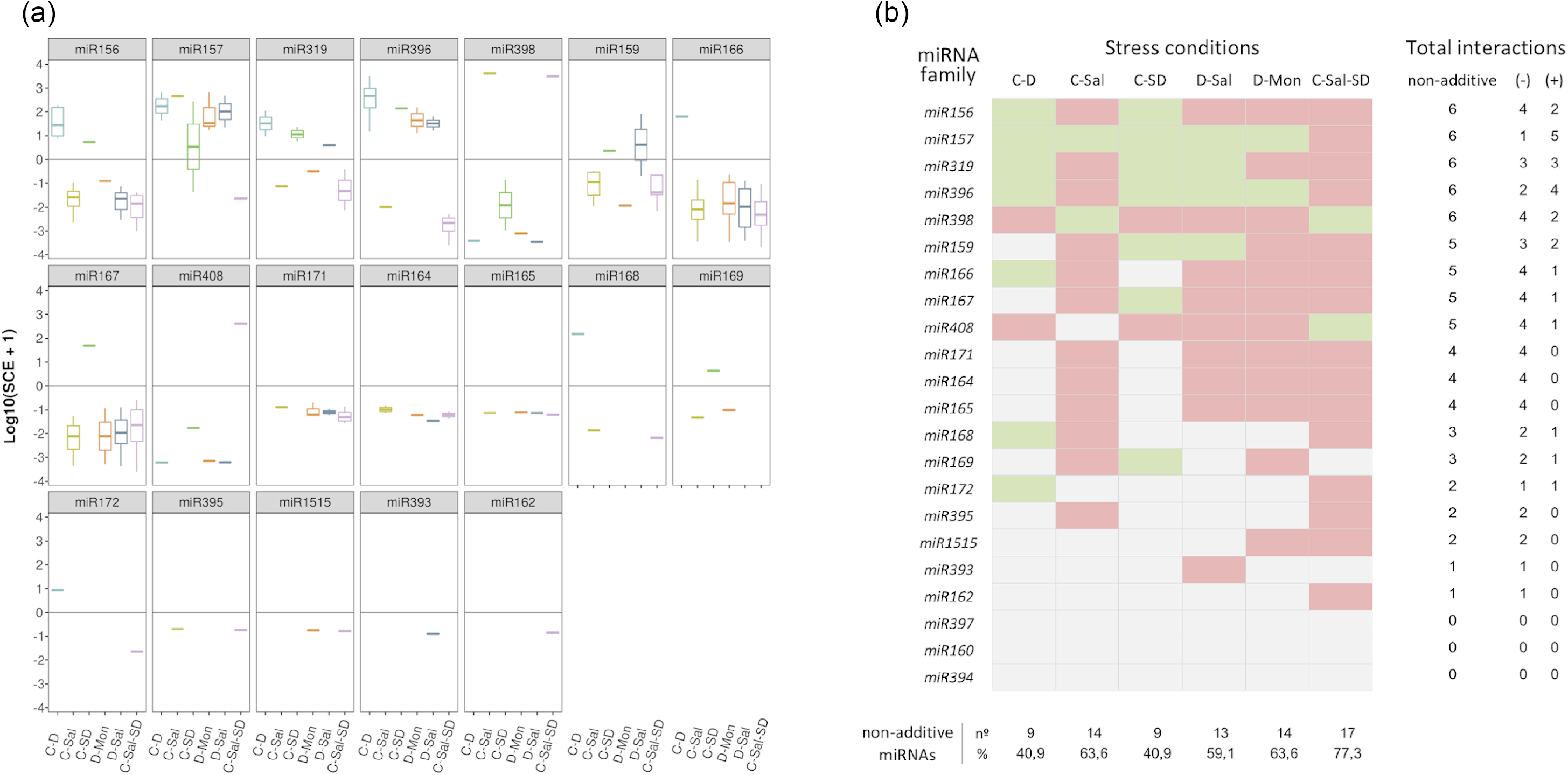
Members of each miRNA family respond in a coordinated manner to combined stresses. (a) Boxplot analysis showing the *SCE* values for family-miRNA related members in each combined stress condition. To determine the general sense of the effect induced by combined stresses for each miRNA family we employed the median of the *SCE* values obtained for the totality of the family members (represented by internal box-line). (b) Graphic representation of the global non-additive positive (green) or negative (red) effects associated to combined stresses estimated for each miRNA family in the six stress conditions analyzed here. The number of combined stresses that induce positive and/or negative non-additive responses in each miRNA family is detailed in the right columns. The proportion of miRNA families with non-additive effects in response to each combined stresses is detailed below.

### 3.4 Different miRNAs families act distinctively in response to combined stresses

To get further insights into the response of each miRNA family to combined stress conditions, we analyzed the rate of differential response to double and triple stresses. The 22 stress-responsive miRNA-families were organized into a table of presence and absence (Table S8) in which the values one and zero represent, respectively, whether or not a miRNA shows a significant response value (with either positive or negative effect) under a combined stress condition. Members of miR156, miR157, miR319, miR396, and miR398 families showed significant positive or negative *SCE* in the six stress conditions analyzed here, while miR159, miR166, miR167, and miR408 members accumulate differentially in five stresses combinations. Sequences belonging to miR164, miR165, miR171, and miR393 (with positive or negative *SCE* in four conditions), miR168 and miR169 (in three), miR172, miR395 and miR1515 (in two) and miR162 (negative effect under C-Sal-SD), showed the lowest differential accumulation in response to the combined stress. Responsive miRNAs included in the miR160, miR394 and miR397 families lacked of significant interactions in any the six analyzed stress conditions.

Correlation between miRNA responses (considering miRNA behavior and the different combined treatments) was estimated by multi-cluster analysis (MCA). MCA evidenced that the response values to combined stresses can be organized into three significantly different groups (Figure 5A). The group including miR156, miR157, miR166, miR319, miR396, miR398, and miR408 contained the miRNA families that exclusively show significant non-additive response values (*SCE* ≠ 0 values) to combined stress conditions. In contrast, families (miR160, miR162, miR168, miR172, miR394, miR397, miR395, and miR1515) with predominantly independent responses were clustered in the second group. Families of miRNAs in which the proportion of significant (*SCE* ≠ 0 values) and non-significant (additive *SCE* values) response was comparable (miR159, miR164, miR165, miR167, miR169, miR171, and miR393) were also clustered together.

**Figure 5.**
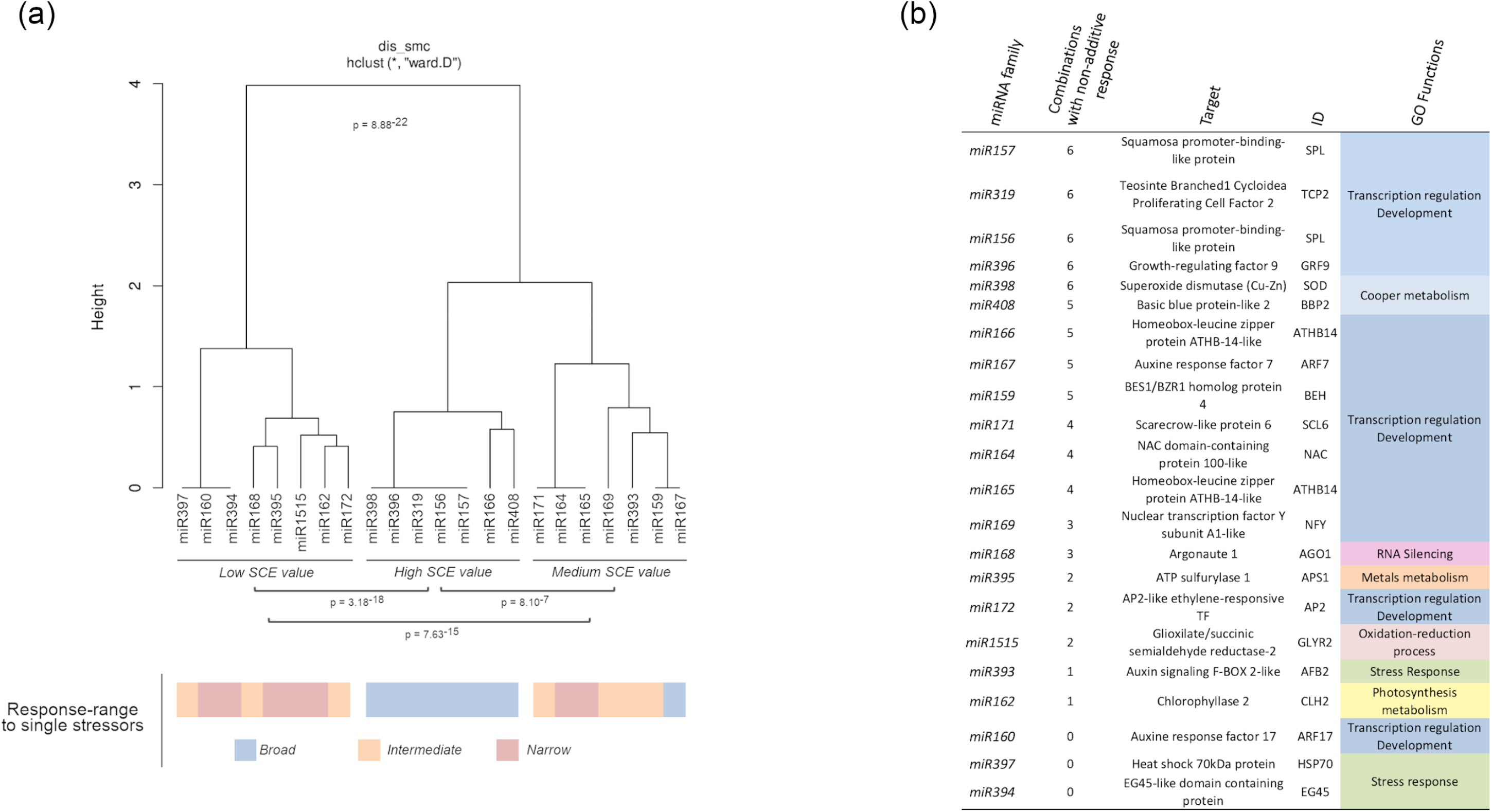
Biological functions of miRNAs with non-additive response to combined stresses. (a) Dendrogram showing the clustering of miRNAs families with at least a member with significant non-additive response to combined stresses in three main groups according to their *SCE* values in the analyzed stress conditions. The global statistical significance of the identified clusters (*p* = 8.88×10^−22^) was estimated by Mann-Whitney-Wilcoxon test, considering the inter- and intra-group Euclidean distances. The lower panel shows the response range determined for each miRNA family in response to single stresses with both biotic and abiotic source (using a color scale). (b) Description and detailed information of the targets for miRNAs with significant non-additive response to combined stresses identified in melon plants. The GO terms were estimated in base to information of homologous transcripts in *A. thaliana*.

Interestingly, all the miRNAs clustered in the group showing significant non-additive expression in response to combined stresses correspond to melon miRNA families already identified as reactive to a broad range of stress (generalists) (Sanz-Carbonell, Marques, Martinez & Gomez, 2020), while miRNAs characterized by a narrow response range (specialists) are the most frequent class (five out eight) in the group showing mainly an additive response to double and triple stresses (Figure 5A - lower part). Finally, miRNAs identified previously as intermediates, are mainly (four out seven) included in the group where significant and non-significant response to the combination of stressor was observed at comparable frequencies. The specialist miRNAs exhibit exclusively *SCE* < 0 response to double and triple stresses, whereas miRNAs identified as generalists showed an even distribution of significant non-additive responses (20 positive and 25 negative *SCE* values). Intermediate miRNAs, although showed a few miRNAs (five) with positive effects, were predominantly (sixteen miRNA families) characterized by a negative response to the combination of stresses. The relationship between miRNA trend response and stress condition was generally dependent of the specific stress/miRNA interaction, although the miR398 and miR408 families showed a coordinated response in all the analyzed conditions, with the exception of C-Sal. However, a positive response (*SCE* = 654.96, *p* = 0.04) was observed for miR408, in this condition, although was considered as non-significant based in the FDR criterion (Table S6). This specifically coordinated activity of the miR398/miR408 tandem was particularly evident in response to, C-SD and C-Sal-SD in which their response was the opposite to the general trend observed for the remaining miRNA families.

Regarding miRNA-regulated targets, it was evident that miRNAs involved in the regulation of transcription factors (TF) associated to plant-development exhibit the higher rate of differential response to combined stress (Figure 4c). In contrast miRNA families expected to modulate the expression of transcripts related (according to GO terms) to a more diverse range of biological functions (RNA silencing, metals metabolism, photosynthesis, response to stress, etc.), showed predominantly a non-significant response to stresses combination.

## 4 DISCUSSION

Much effort has been dedicated to elucidating the mechanisms underlying stress response in crops. Although great progress has been made in the last years, including the identification of both protein-coding and non-coding transcripts responsive to different stresses, most studies focused on deciphering the plant regulatory pathways triggered in response to single stress conditions. Alas, no much effort has been devoted to understand the plant responses to multiple stresses acting simultaneously; a situation that is most common in the wild.

Here, we have addressed this question by measuring the miRNA-mediated responses to combined stresses in melon plants exposed to five different double and one triple stressful condition. Our strategy comprises two principal steps, first to identify the miRNA-families responding to double and triple stress conditions. Second, we compared the expression level of such responding miRNAs with the values previously obtained in melon plants exposed to the respective single stresses. This comparative analysis has allowed us to determine how the stress combinations affect the differential expression of miRNAs; disentangling stress-specific responses to general responses. This information enabled the inference of the global structure of the miRNA-mediated differential response to combined stress conditions in melon.

The computational analysis identified 22 miRNA families with significant differential expression in response to the analyzed stresses. Regarding their functional role, these reactive families mainly target melon homologous to well-described TFs (e.g., SPOROCYTELESS, BES1/BZR1 HOMOLOG 4, AUXINE RESPONSE FACTORS (ARF), ARABIDOPSIS THALIANA HOMEOBOX PROTEIN 14, TEOSINTE BRANCHED 1/CYCLOIDEA/PROLOFERATING CELL FACTOR, APETALA 2, GENERAL REGULATORY FACTOR (GR), and NUCLER FACTOR Y). This is in agreement with previous observations in other species (*A. thaliana*, rice, maize, sorghum, sunflower, etc.) in which it has been reported that in general, miRNAs reactive to stress target predominantly TFs (Samad et al., 2017). This reinforces the emerging notion that the role-played by miRNAs during the stress response is evolutionary conserved in plants (Rubio-Somoza & Weigel, 2011; Megraw, Cumbie, Ivanchenko & Filichkin, 2016; Sanz-Carbonell, Marques, Martinez & Gomez, 2020) and emphasizes the potential of miRNAs as targets for improving stress tolerance in crops (Tang & Chu, 2017; Chaudhary, Grover & Sharma, 2021). The totality of these stress-responsive miRNA families were coincident with the previously described as reactive in single biotic and abiotic stress conditions in melon (Sanz-Carbonell et al. 2019; Sanz-Carbonell, Marques, Martinez & Gomez, 2020). The observation that double and triple stresses do not induce the differential accumulation of any miRNA family reactive specifically to combined stress, suggest that (at least under the conditions analyzed here), the miRNA families involved in the response to stress comprise the general structure that modulate the recovery of the plan-cell homeostasis under both single and combined adverse environmental conditions.

Considering the response rate to each stress-combination we observed a more consistent activity in certain miRNA families. Our results evidenced, that melon miRNAs (miR156, miR157, miR166, miR167, miR319, miR396, miR398, and miR408) previously characterized by exhibit differential accumulation in response to a wide range of biotic and abiotic stress conditions in melon, maize and soybean (dubbed as generalists), were differentially expressed in the six analyzed conditions, evidencing a high response range, independently of the stresses combination. Interestingly, miRNAs families reactive to four or less conditions (miR162, miR164, miR165, miR172, miR394, miR397, miR395, and miR1515) predominantly corresponded to miRNAs characterized by exhibiting differential response to specific stresses (specialists). It has been recently suggested that generalists stress-responsive miRNAs might be involved in the modulation of the central steps in the recovery of the cell homeostasis during the exposition to adverse environmental conditions, while specialists families responding to specific stress conditions and/or exposition times had been hypothesized to be involved in the regulation of metabolic processes associated to each particular stressor (Sanz-Carbonell *et al*. 2019; Sanz-Carbonell, Marques, Martinez & Gomez, 2020). Assuming this responsive behavior, it is expected that generalist miRNAs were the predominant class reactive to double and triple stresses. Sequences related to generalist miRNA-families are characterized by mainly modulating master regulators or central hubs, predominantly TFs related with plant development (Sanz-Carbonell, Marques, Martinez & Gomez, 2020). It is well established that alteration in the expression of TF genes normally results in remarkable changes in the global gene expression during plant growth and development (Li et al., 2015). Furthermore, it has been proposed that such TFs might, for example by co-regulatory feedback and feedforward loops miRNA/TF, act as amplifiers of the plant-response to stress (Rubio-Somoza & Weigel 2011; Megraw et al., 2016; Samad et al., 2017). The generalist class is comprised by miRNAs previously described as reactive to different biotic and/or abiotic stress conditions in diverse plant-species. Several studies support that the module miR156-SPLs besides exhibiting a broad response range to low temperatures in diverse plant-species (Zhou & Tang, 2019), also improves tolerance to salinity, heat and drought in *Medicago sativa* (Arshad, et al., 2017; Arshad, Gruber, Wall & Hannoufa, 2017; Matthews, Arshad & Hannoufa, 2019). Moreover, the interaction between miR396 and GRF is involved in the modulation of the response to diverse biotic (*Phytophthora nicotianae*) and abiotic (drought, salt, alkali, UV-B radiation, and osmotic unbalance) stress conditions (Gao et al., 2010; Kim et al., 2012; Casadevall et al., 2013; Chen, Luan & Zhai, 2015). Cotton plants overexpressing miR157 suppressed the auxin signal and showed enhanced sensitivity to heat (Ding et al., 2017). Recent studies evidenced a critical function for miR166 in tolerance to abiotic stresses in maize (Li et al., 2020) and cadmium-induced toxicity in rice (Ding et al., 2018). By means of transgenic approaches it was established that miR167 acts as transcriptional regulator in response to bacterial infection (Jodder, Basak, Das & Kundu, 2017) and temperature-induced stress in tomato plants (Jodder et al., 2018). Multiple evidences obtained by both sRNA-sequencing and transgenic approaches, support the role of members of the miR319-family, an ancient miRNA conserved across plant species ranging from mosses to higher plants, as a key modulator of the plant-environment interrelation (at biotic and abiotic level) in monocotyledonous and dicotyledonous species (Bustamante et al., 2018; Liu et al., 2019; Shi et al., 2019; Wu, Qi, Meng & Jin, 2020; Fang et al., 2021; Joshi, Chauhan & Das, 2021). Finally, regarding miR398 and miR408 families, it was recently proposed that these conserved miRNAs, involved in the maintenance of the cooper homeostasis in plants, might be also involved in the systemic signaling of the response to biotic and abiotic stresses (Burkhead et al., 2009; Sanz-Carbonell, Marques, Martinez & Gomez, 2020).

Upon determining the melon miRNAs responsive to combined stress conditions, we attempted to analyze whether the expression of these stress-responsive miRNAs was different in comparison with that observed under each one of the stresses individually. Our conceptual premise assumes that miRNAs that did not show a significant differential (positive or negative) response to combined stresses exhibit an independent behavior to the combination of the stress conditions. The obtained results demonstrated that in a considerable proportion of the analyzed miRNA-stress combinations (59.85%), the stress-responsive miRNAs families exhibit a differential response to the action of combined stresses. This evidences that, although the miRNAs involved in the regulation of the response to a particular stress combination are coincident with such described under individual stresses, the regulatory effects exerted on their targets is considerably different when the plant is exposed to a combination of adverse environmental conditions.

Considering in detail the differentially reactive miRNAs, we observed that generalist miRNAs showed the higher rate of differential accumulation (compared to the observed respect the response to single stresses) in response to combined adverse environmental conditions. Thus, supporting that the biosynthesis and/or processing of such miRNA-families is particularly (and differentially) susceptible to the combined exposition to two or three stress conditions. In contrast, the data obtained when miRNAs identified previously as specialists were analyzed evidenced that the expression of this class de miRNA families is predominantly independent of the effects of the combined-stresses and corresponds principally to the expression levels observed in response to each stressor individually. This functional behavior of responsive miRNAs to combined stresses is compatible with the architecture of the miRNA-mediated regulatory network of response to adverse environmental stimuli described recently in melon (Sanz-Carbonell *et al*. 2019; Sanz-Carbonell, Marques, Martinez & Gomez, 2020). Structurally, this network is characterized by exhibiting a central core of highly connected miRNAs (generalist), and another peripheral layer comprised of miRNA families with lower connectivity (*specialists*) (Figure 6A). According to this structure, it is expected that the expression of generalist miRNAs (highly interconnected and reactive to a broad range of stress conditions) might be differentially affected (either positively or negatively) by the incidence of two or more distinct stresses (Figure 6B). In contrast, specialist miRNA-families (with low connectivity and reactive to particular stress conditions) remain functionally independent to the effects of additional non-related stresses, and respond mainly to the exposition to combined stress conditions in additive (non-differential) manner (Figure 6A). The observation that the architecture of the miRNA-mediated regulatory network of response to stress in melon is able to predict the predominant reactivity rate of the miRNA-response to combined stresses, provide additional robustness to this inferred regulatory structure involved in the miRNA-mediated modulation of plant-environment interactions. Furthermore, the fact that a structurally comparable miRNA-networks of response to stress has been also proposed in rice and soybean plants exposed to diverse biotic and abiotic stress conditions (Sanz-Carbonell, Marques, Martinez & Gomez, 2020), allows to speculate about the possibility that the response pattern to combined stresses observed in melon may well be extended to another crops.

**Figure 6.**
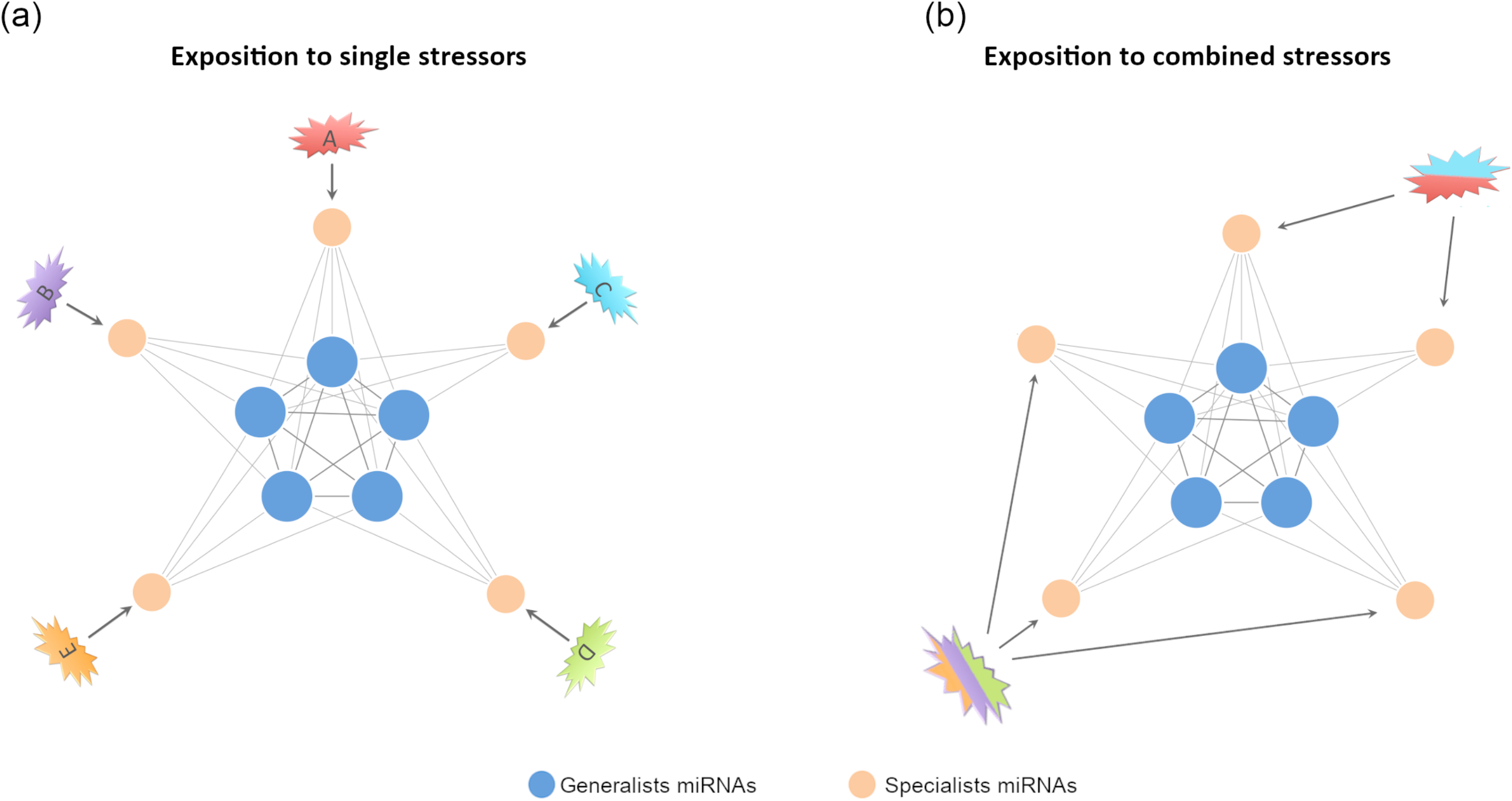
Proposed model to explain predominant non-additive response in certain miRNAs families. (a) Simplified graphic representation of the proposed miRNA-mediated network of response to stress in melon (Sanz-Carbonell et al., 2019; Sanz-Carbonell, Marques, Martinez & Gomez, 2020). Blue nodes represent highly connected miRNAs with a broad response range to biotic and/or abiotic stress conditions (*generalists*). Orange nodes represent miRNAs reactive to specific stress conditions (*specialists*). (b) When the network is exposed to double or triple stress conditions is expected that the stresses combinations should not affect specialist miRNAs (poorly connected between them) and consequently they exhibit additive *SCE* values (comparable to the resultant of the sum of both individual responses). In contrast, generalist miRNAs (highly interconnected) respond to stresses combination in a differential (non-additive) manner, related to each stress combination.

In general, the transcripts of well-established TFs were the targets modulated by miRNAs with significant non-additive effects in response to combined stresses, reinforcing the key role assumed for the circuits miRNA-TF in the regulation of the stress response in plants (Rubio-Somoza & Weigel, 2011). Regarding the trend of the global differential miRNA-mediated response to combined stresses negative values were the most abundant. Response values lower than the expected for stress-independent effects might be initially assumed as an indicative of functional convergence in the miRNA-mediated response to combined stresses. It has been recently suggested that specific developmental events may be usually modulated by diverse miRNAs in rice (Tang & Chu, 2017). In this proposed model, miRNAs functionally converged via direct or indirect interaction between their targets. It is well established that osa-miR393 regulate the auxin receptors OsTIR1 and OsAFB2, both involved in the ubiquitin-mediated degradation of specific substrates during auxin signaling (Bian et al., 2012; Li et al., 2016). Furthermore, osa-miR160 and osa-miR167 modulate the expression of at least three ARF transcripts (OsARF8, OsARF16 and OsARF18) (Yang, Han, Yoon & Lee, 2006; Li et al., 2014; Huang, Li & Zhao, 2016). Interestingly, cmel-miR393 and cmel-miR167 exhibit a predominant negative differential response (assumed as indicator of functional convergence) to the combined stresses analyzed here. Further studies are needed to determine the existence of a potential functional convergence in the miRNA-mediated response to multiple stresses.

Altogether, our results provide additional support to the anticipated notion that plants may use the miRNA-mediated regulation as pivotal mechanism to recover the cell homeostasis in response to both simple and combined stresses (Zhang, 2015; Samad et al., 2017; Zhu et al., 2019; Zhou et al., 2020). The confirmation that the previously described as generalist miRNAs are also the predominant components of the global miRNA-mediated response to combined stress conditions highlights the possibility that this class de miRNAs may emerge as a valuable breeding-target for improving, in the near future, crop tolerance to the multiple adverse environmental conditions associated to climate change.

## ACKNOWLEDGEMENTS

J.M.M. is recipient of a predoctoral contract ACIF-2017-114 from the Generalitat Valenciana.

## AUTHOR CONTRIBUTIONS

PVB: Performed and designed computational analysis, prepared figures, and discussed the results. JMM: Analyzed the results, prepared figures and contributed to write the manuscript. MCM: Conceived and performed RT-qPCR analyses and discussed the results. AGHA: performed RT-qPCR analysis. JCS: Performed computational analysis. BP: Provide the *Monosporascus* isolate and contributed to design the stress treatments. AJM: Provided melon seeds and contributed to design the stress treatments. SFE: Conceived and perform the estimation of the *SCE* values and revised the manuscript. GGG: Conceived and designed the experiments, analyze the results and drafted the manuscript. Manuscript review: All authors read and approved the final manuscript.

---

**Figure S1.**
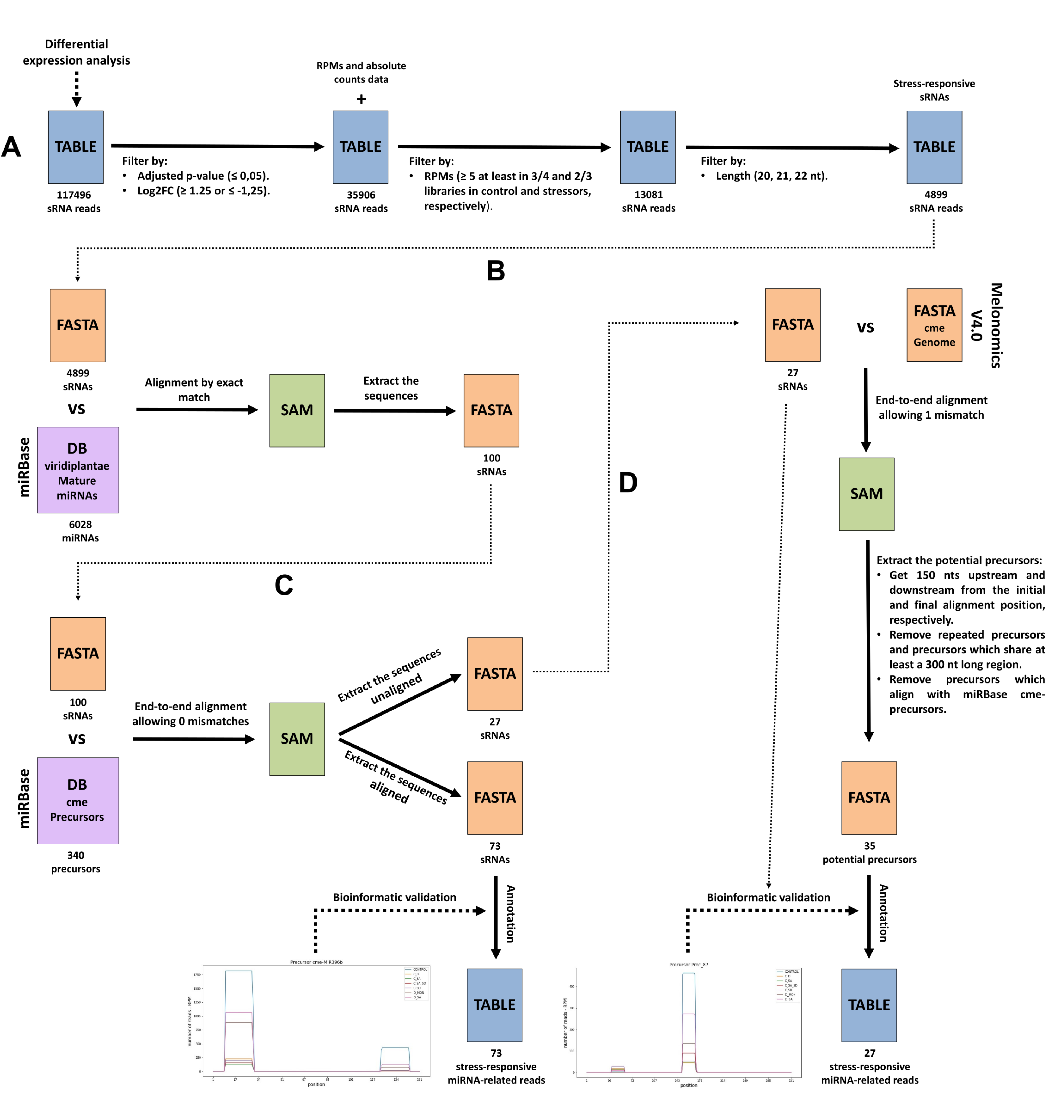
Pipeline for miRNA detection. **A** sRNA reads data coming from the differential expression analysis were filtered by adjusted p-value, log2FoldChange, RPMs and length to get the stress-responsive sRNA reads; **B** Stress-responsive sRNAs were aligned by exact match on viridiplantae mature miRNAs deposited in miRBase; **C** Stress-responsive sRNAs matched in the previous step were aligned on the cucumis melo precursors deposited in miRBase without allowing mismatches. The aligned sequences were bioinformatically validated and annotated as miRNA to be used in this work; **D** Stress-responsive sRNAs unaligned in the previous step were aligned on the cucumis melo genome regarding biological variability, that is, allowing 1 mismatch. Then, we looked for potential precursors which were used to bioinfomatically validate the sequences as miRNA. These sequences were annotated taking into account only the miRNA family of the viridiplantae mature miRNA on which aligned in the step B.

**Figure S2:**
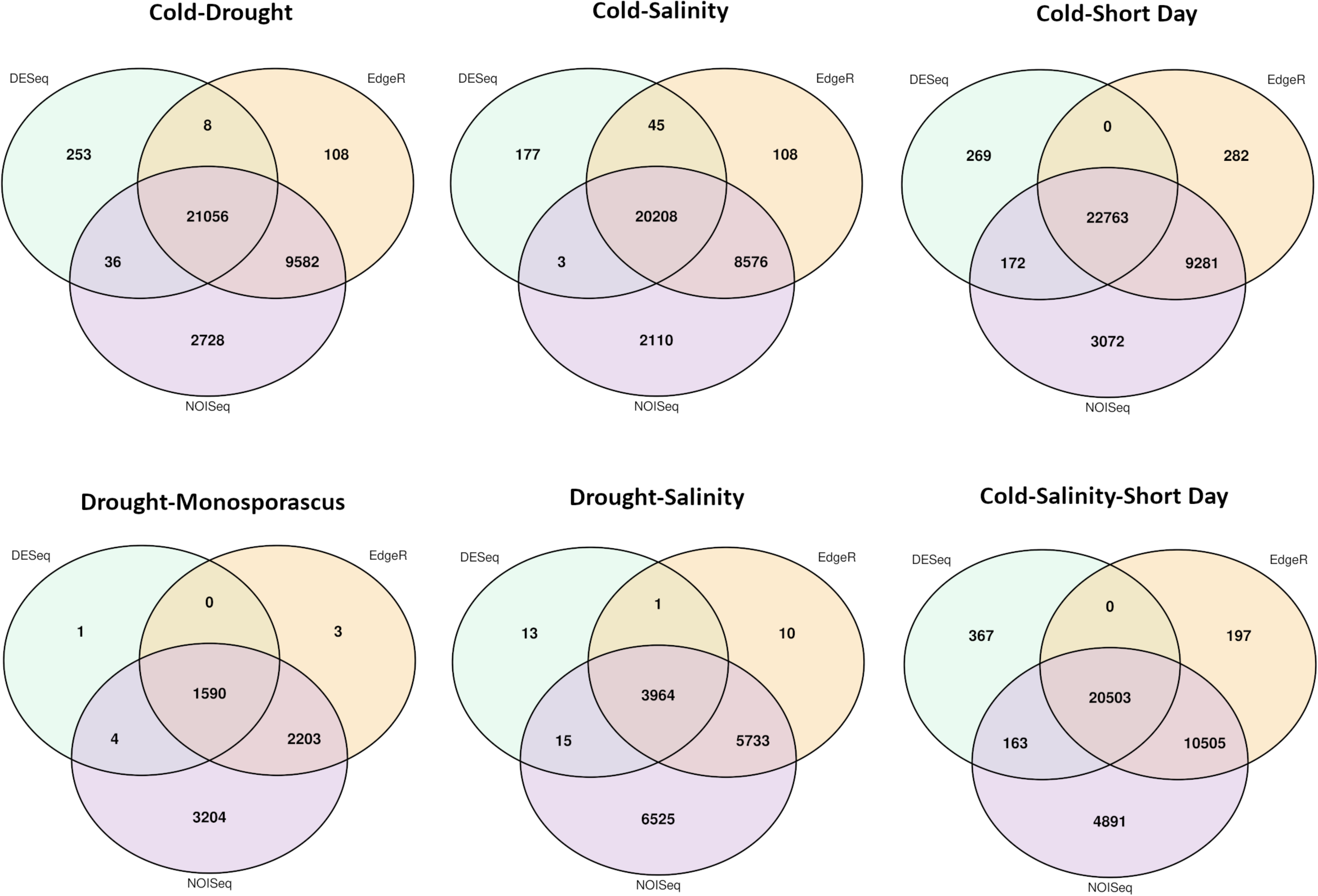
Analysis of stress-responsive miRNAs. Venn diagram comparing the number of the differential sRNAs - estimated by DESeq2 (green), edgeR (orange) and NOISeq (magenta)- expressed in melon in response to combined stress conditions. Only the sRNAs predicted as differential by all three analysis methods were considered as true stress-responsive miRNAs.

**Table S1:** Detail of the combined stress tratments used in this work.

| Stress Conditions | Treatments at 11 days post emergency |  | sample collection |
| --- | --- | --- | --- |
| *Cold Drought* | Irrigated with 50 ml Hoagland's solution<br>20 °C/16 h-light --- 14 °C/8 h-darkness | Except for drought combined treatments plants were irrigated alternatively (water and Hoagland's solution) by inundation (1500 mL / 48 hs) | 11 days after stress treatment |
| *Drought Salinity* | Irrigated with 50 ml of LiCl (200 mM)<br>28 °C/16 h-light --- 20 °C/8 h-darkness |  |  |
| *Cold Salinity* | Irrigated with 50 ml of LiCl (200 mM)<br>20 °C/16 h-light --- 14 °C/8 h-darkness |  |  |
| *Cold Short-day* | Irrigated with 50 ml Hoagland's solution<br>20 °C/8 h-light --- 14 °C/16 h-darkness |  |  |
| *Drought Monosporascus* | Irrigated with 50 ml Hoagland's solution plus *M. cannonballus* mycelium (1000 UFC)<br>28 °C/16 h-light --- 20 °C/8 h-darkness |  |  |
| *Cold Salinity Short-day* | Irrigated with 50 ml of LiCl (200 mM)<br>20 °C/8 h-light --- 14 °C/16 h-darkness |  |  |
| *Control* | Irrigated with 50 ml Hoagland's solution<br>28 °C/16 h-light --- 20 °C/8 h-darkness |  |  |

**Table S2:**
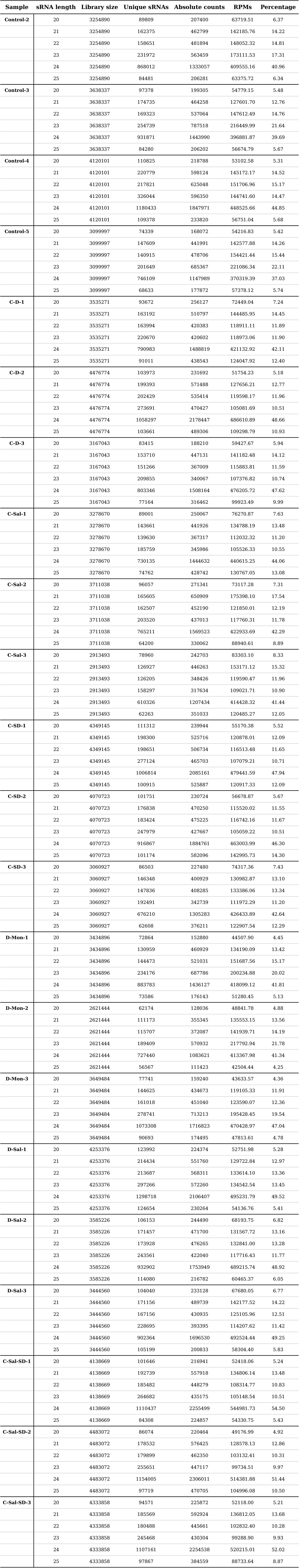
Detailed information of control and stress combined libraries of Cucumis melo by sRNA length.

**Table S3:**
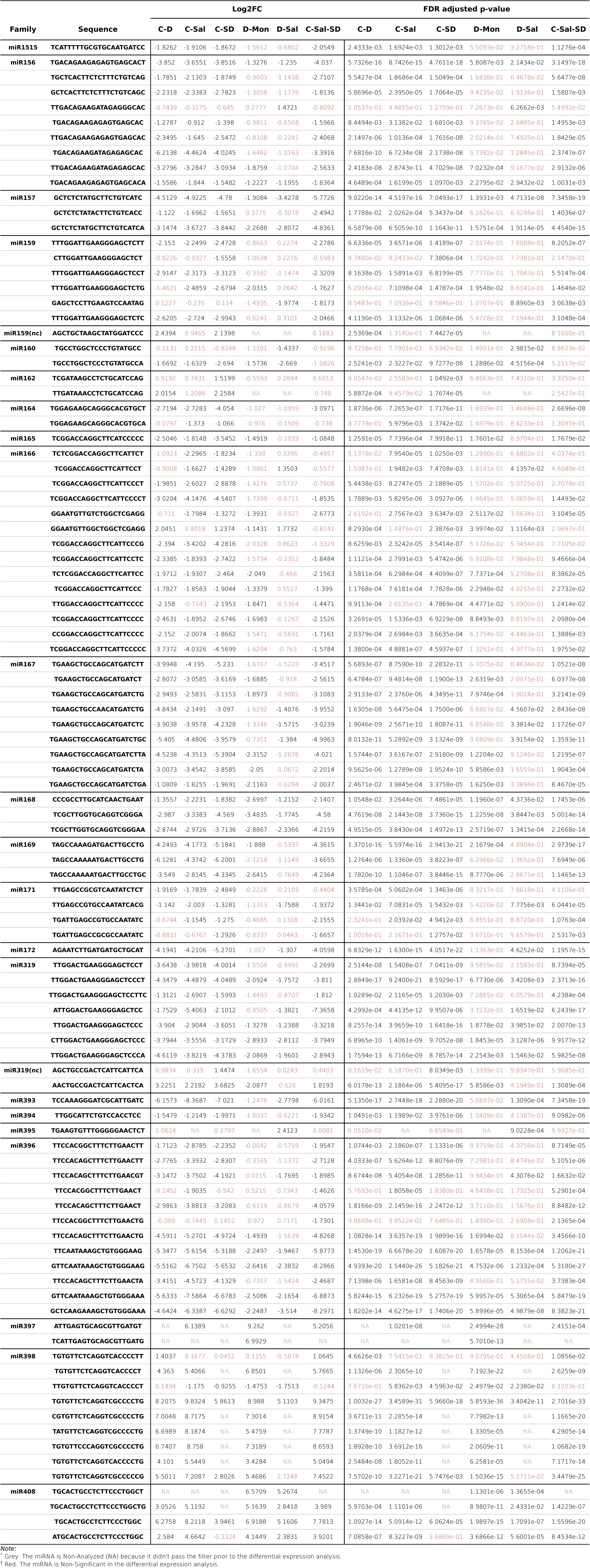
Log2FC values and FDR adjusted p-values of stress-responsive cucumis melo miRNAs in stress combined conditions.

**Table S3a:**
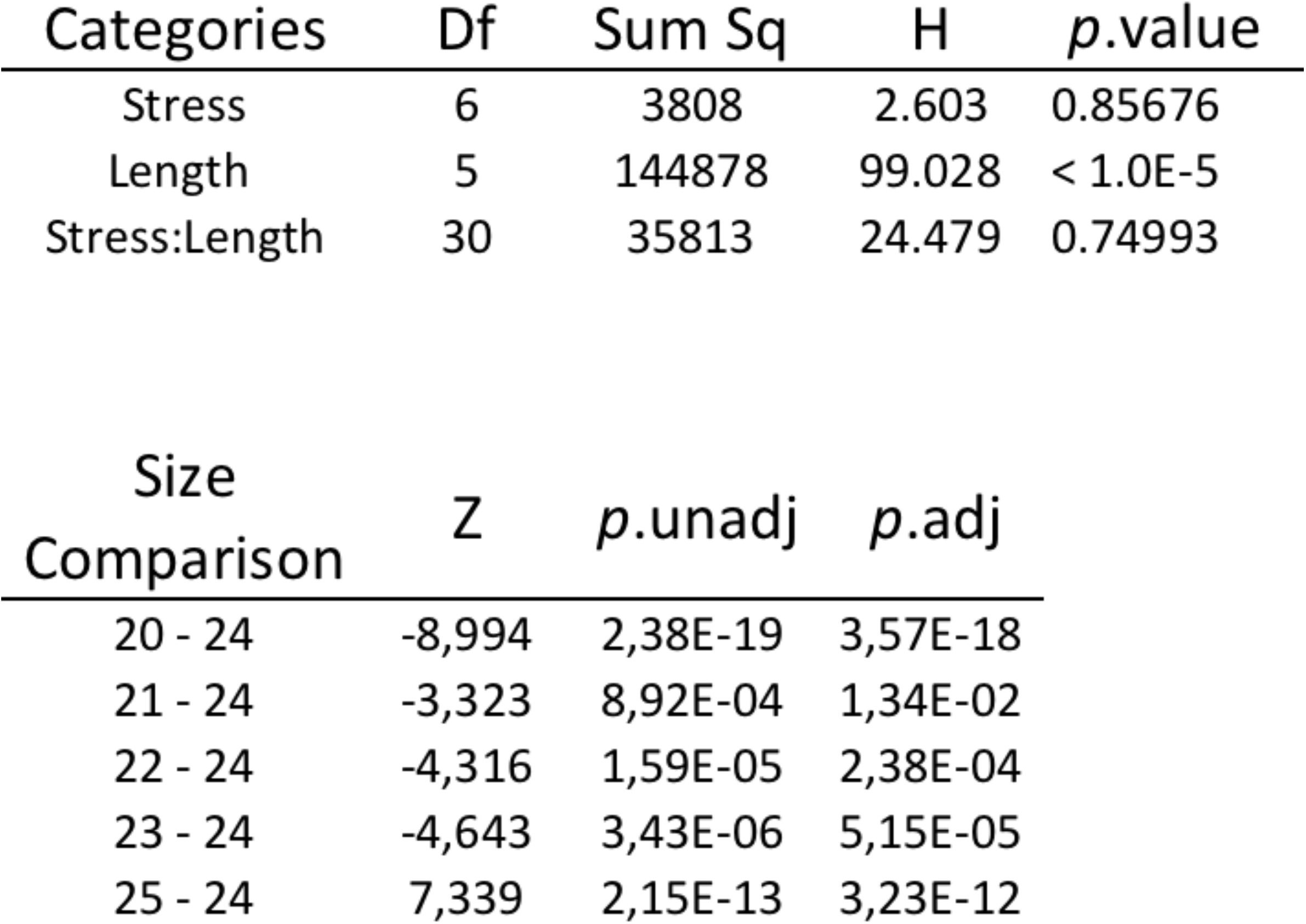
Statically analysis of sRNAs-reads profiles in control and stresses exposed plants. The differences between treatment and reads-length were analyzed by the Scheirer–Ray–Hare non-parametric test (upper). Once established that only length category shown significant alterations we used Dunn’s Multiple Comparison Test to analyze the difference between 24 nt length reads and the rest of the read-size categories (lower).

**Table S4:**
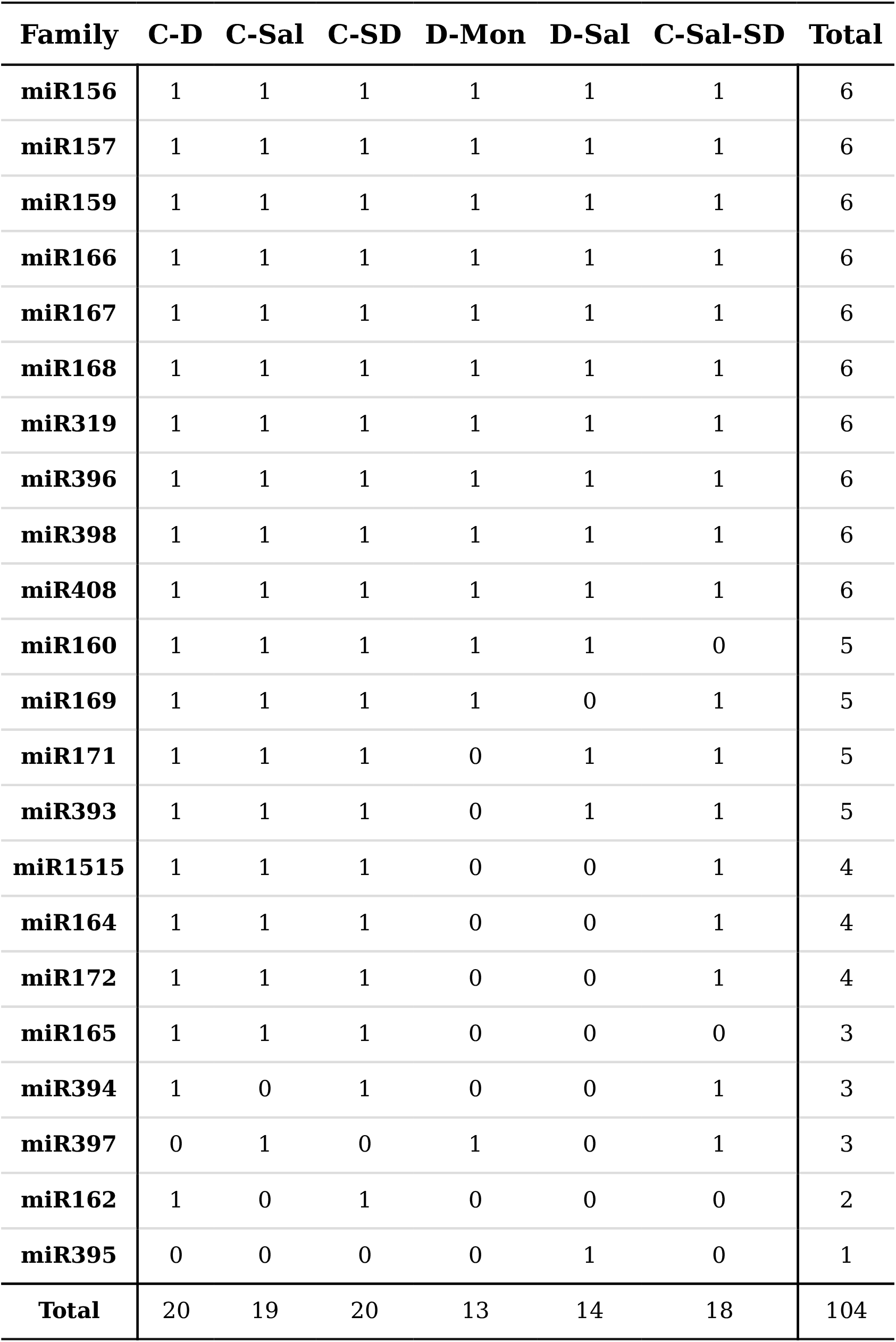
Presence and absence of stress-responsive miRNAs to a combined stress conditions in Cucumis melo. 1: stress-responsive, O: non stress-responsive.

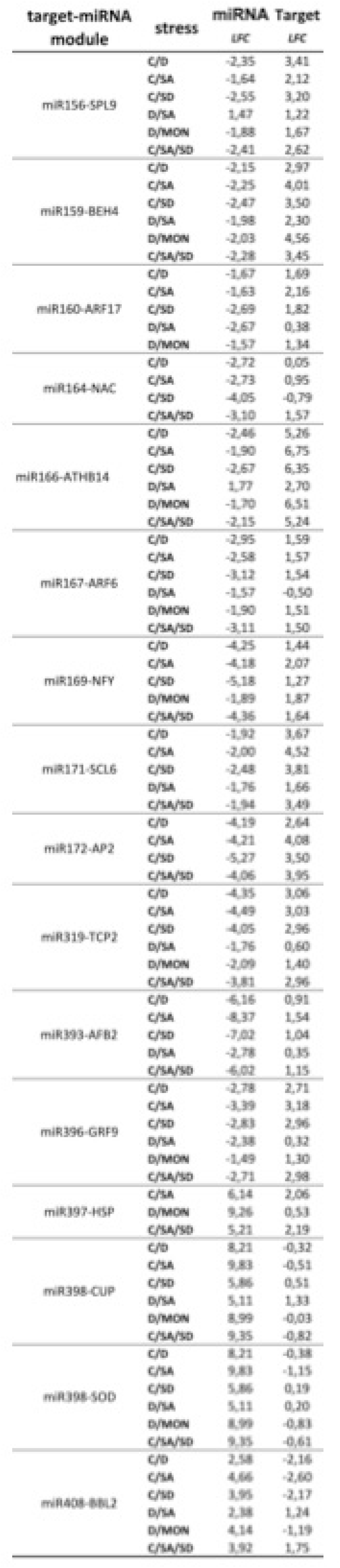

**Table S6B:**
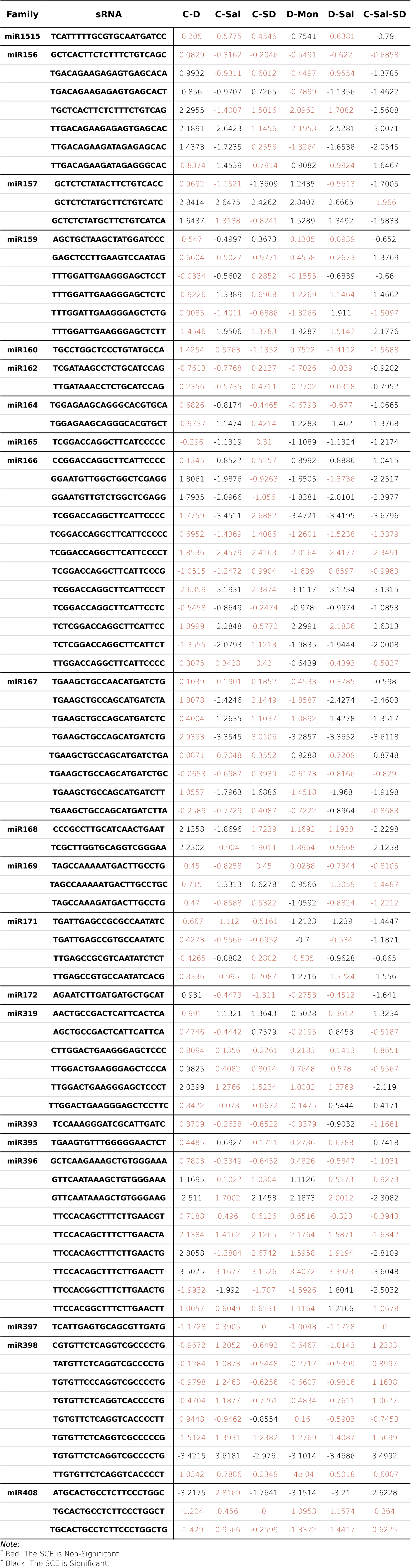
Log10(SCE + 1) values of stress-responsive miRNAs in Cucumis melo.

**Table S7:**
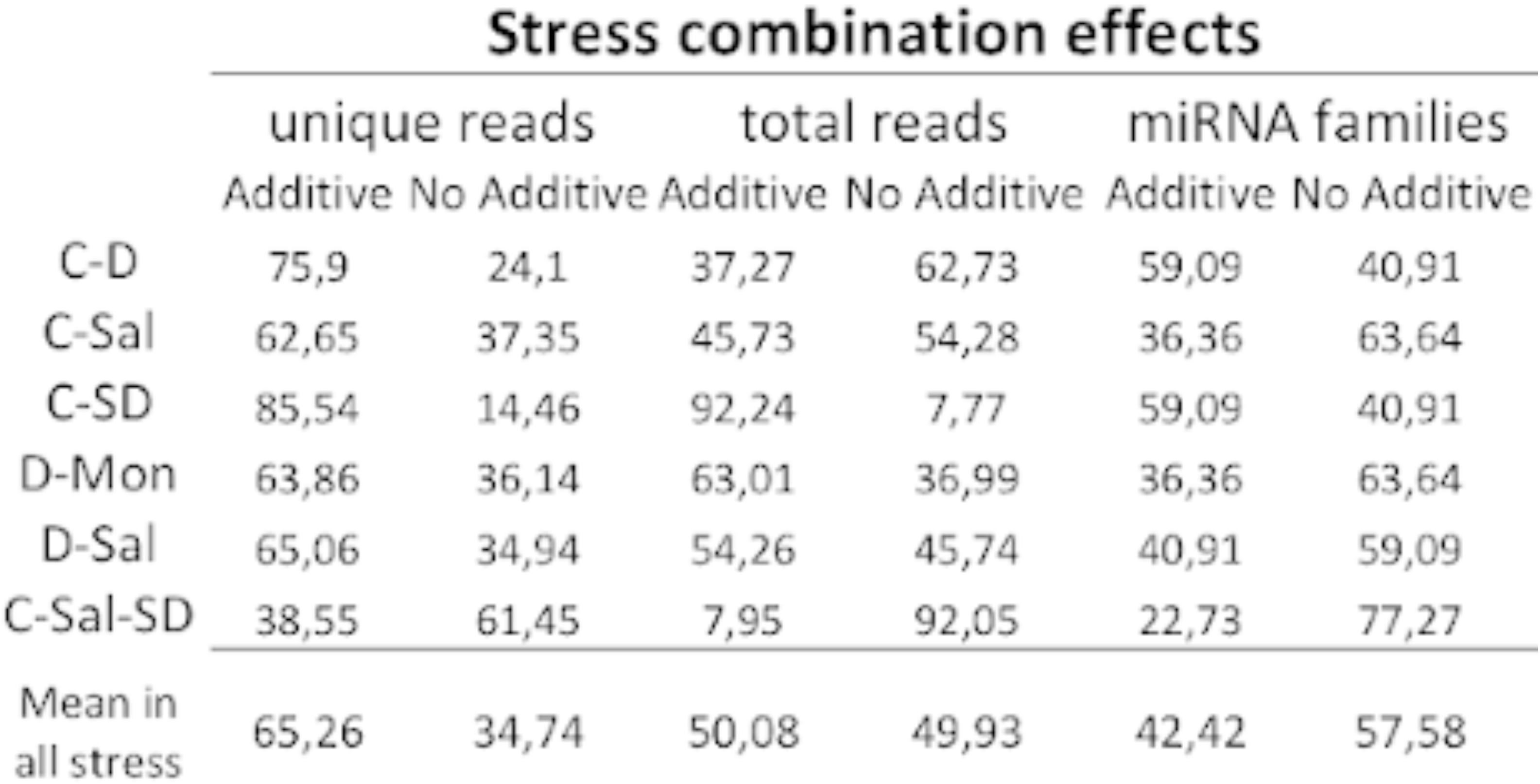
Detail of the percentage of additive and non-additive values SCE values obtained for differentially expressed miRNAs in each analyzed stress combination.

**Table S7:**
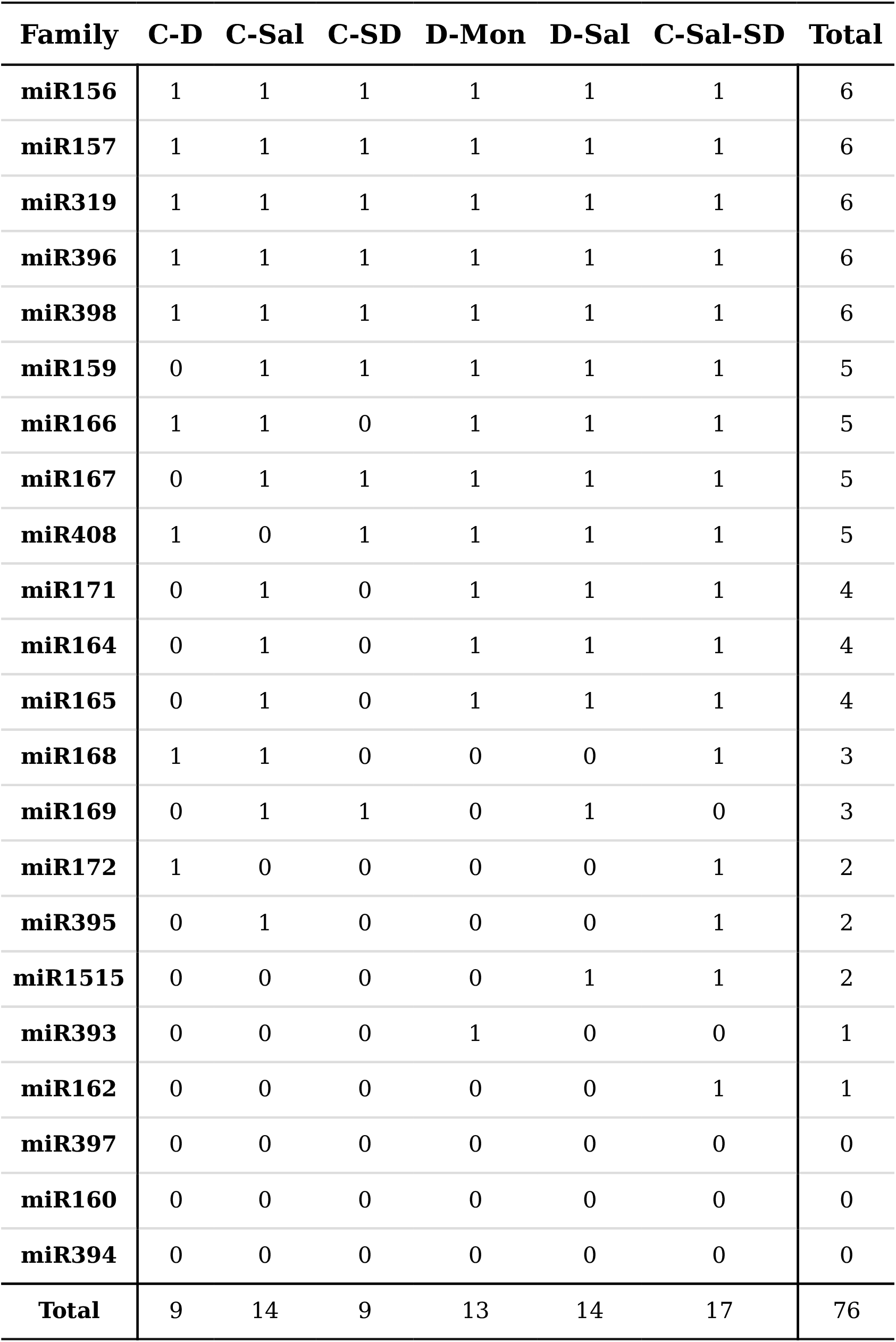
Presence and absence of Stress Combination Effect (SCE) for stress­ responsive miRNAs in Cucumis melo. 1: non-additive SCE, O: additive SCE.

